# Reduced PDE4D7 expression in prostate cancer correlates with down-regulation of genomic elements within the up-stream PDE4D coding region on chromosome 5

**DOI:** 10.1101/2022.06.16.496387

**Authors:** Chloe Gulliver, George S Baillie, Ralf Hoffmann

## Abstract

PDE4D7 expression is diminished during progression of PCa and the phosphodiesterase has been proposed as a prognostic biomarker. RNA sequencing of PCa tissue identified sequences in the PDE4D coding region on Chr5q12 exhibiting similarities in mRNA expression profile to PDE4D7. As previously identified, PART1 had a significantly similar expression pattern to PDE4D7 across samples. However, other genes also matched expression to PDE4D7, including miRNAs and lncRNAs. These novel PDE4D7 associated genes represent putative PCa biomarkers and could have mechanistic roles in PCa progression.

## Introduction

Given the substantial heterogeneity in prostate cancer (PCa) treatment responses vary, therefore a prognostic biomarker indicating disease progression would influence personalised treatment decisions and identify novel therapeutic targets (1). Dysregulated cyclic AMP (cAMP) signalling is associated with cancer, particularly in the progression from localised androgen-sensitive (AS) PCa to aggressive castration-resistant PCa (CRPC) (2). cAMP signalling depends on the compartmentalisation of cAMP effector proteins and phosphodiesterases (PDEs) (3). The PDE4D sub-family is germane in PCa, with the isoform PDE4D7 being of unique importance (4). Interestingly, PDE4D7 expression correlates inversely with disease progression, with downregulation conferring an increased risk of disease recurrence (1,4). This highlights the potential of PDE4D7 as a prognostic biomarker to distinguish between insignificant and aggressive tumours, as well as representing potential novel therapeutic avenues.

## Methods

Two independent NGS transcriptomics expression data sets (n=533 and n=151) from PCa patient biopsy punch samples of surgically resected tumour tissue were used in this study. RNA sequencing and data processing was performed as previously described by van Strijp *et al*. (2019) (5). Generation of normalized PDE4D transcript expression was performed by subtracting the RT-qPCR Cq of respective transcripts from the averaged reference gene RT-qPCR Cq. Normalised PDE4D5, PDE4D7, and PDE4D9 expression was transformed to the PDE4D5, PDE4D7, and PDE4D9 scores (6). To investigate the expression of genomic elements like miRNAs or pseudogenes on chromosome 5 in the region where PDE4D7 exons 1-3 are located (chr5:59,000,000 to chr5:60,500,000), we created a heatmap of the expression of respective transcripts of these genomic elements together with the level of PDE4D5, PDE4D7 and PDE4D9 expression. For this we transferred the PDE4D5, PDE4D7 and PDE4D9 scores as well as the transcript per million (TPM) expression values for the respective transcripts into z-scores on the gene level across all samples. As a result, all genes of the heatmap have a mean expression of 0 and a SD of 1.

## Results

Our analysis suggests a distinct correlation between PDE4D7 expression and that of a subset of lincRNAs, miRNAs and pseudogenes located between 60,000,000 and 60,500,000 base pairs on chromosome 5 [Fig. 1B]. Importantly, all trends observed are correlated between both independent datasets [Figs. 1A (i) and (ii)].

**Figure 1:**
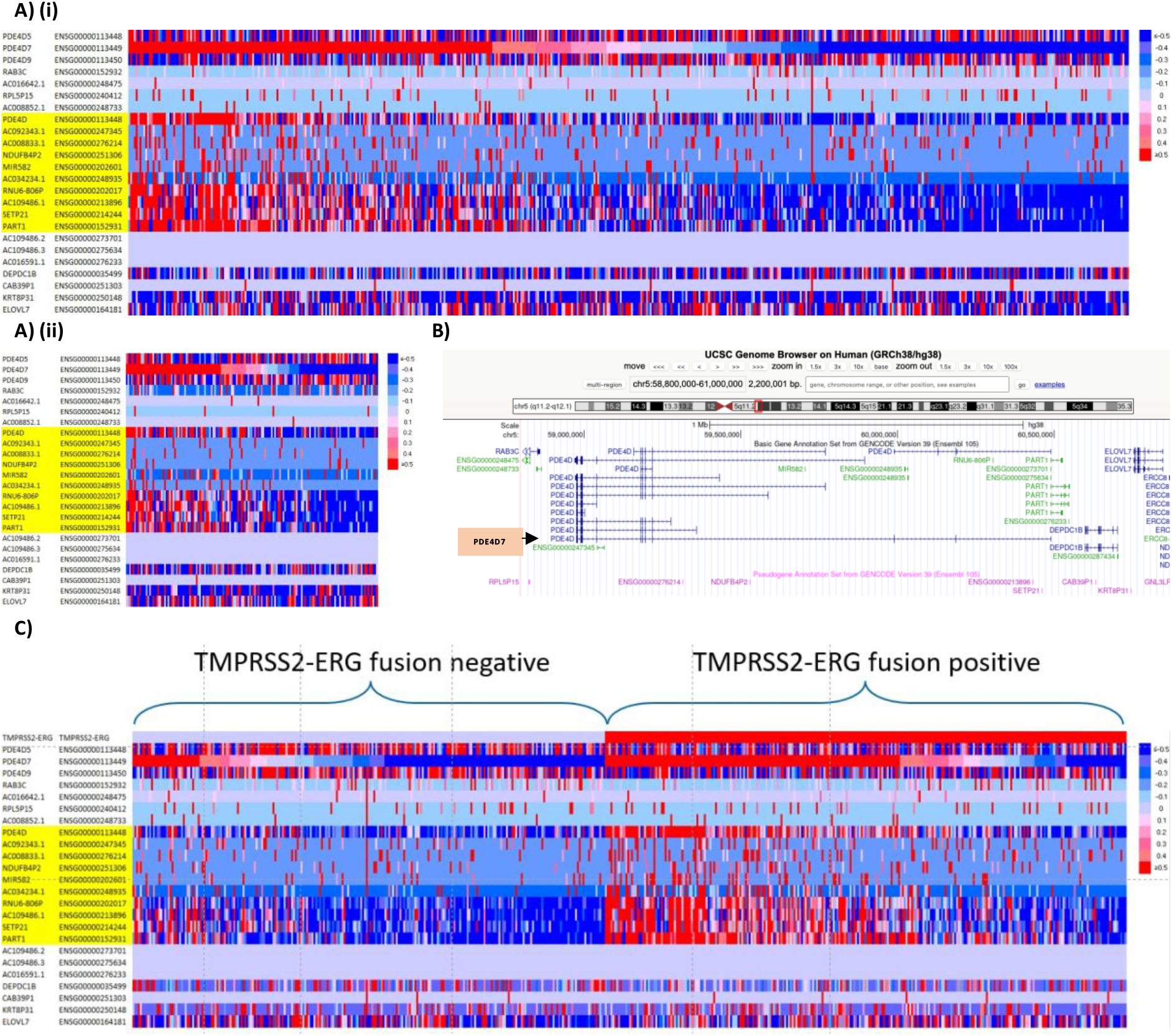
Genome expression on chromosome 5 within PDE4D coding region in prostate cancer patients in comparison to the PDE4D7 score. A) Heat map of gene expression in PCa patient samples from datasets 1 n=533 (i) and 2 n=151 (ii) ordered according to their PDE4D7 scores from high to low. Gene expression values of the shown transcripts located in the genomic region of the PDE4D gene on chromosome 5 were determined by NGS RNA sequencing. Transcript names highlighted in yellow show p-value <0.05 in expression difference between samples with high vs low PDE4D7 score. B) Chromosome 5 genome alignment in PDE4D coding region between approx. 59,000,000 – 60,500,000bp from UCSC genome browser (GRCh38/hg38 Human Dec 2013 assembly), accessed on 14 April 2022. PDE4D7 isoform annotated by arrow and orange text box. C) Heat map of gene expression in PCa patient samples from datasets 1 (n=533) ordered according to their TMPRSS2-ERG fusion status.

Specifically, RNA transcripts located within the first 3 exons of PDE4D7 are coincidentally downregulated, with AC034234.1, RNU6-806P, AC109486.1, SETP21, PART1 and the entire PDE4D gene exhibiting a similar pattern of expression to PDE4D7 among samples. These genes all showed a p-value <0.05 between samples with high vs low PDE4D7 scores [Figs. 1A (i) and (ii)], alongside AC092343.1, AC008833.1, NDUFB4P2 and MIR582.

Whilst AC109486.1 exhibits this pattern, AC109486.2 and AC109486.3 show no expression in any sample, alongside AC016591.1. Interestingly, all samples with no expression are found in the region overlapping PART1 which exhibits similar expression to PDE4D7, revealing that this entire region is not deleted.

CAB39P1 showed almost a uniform lack of expression across all samples, however a minimal number of samples exhibited increased expression irrelevant to PDE4D7 level. AC016642.1, RAB3C, RPL5P15 and AC008852.1 revealed similar lack of expression but with more anomalous samples exhibiting altered expression. The most varied expression profiles were DEPDC1B, KRT8P31 and ELOVL7, as well as PDE4D5 and PDE4D9, reflecting no obvious correlation to PDE4D7 level. Of note here is that TMPRSS2-ERG fusion negative (TMPRSS2-ERG-) patient samples show more pronounced downregulation of genomic elements than TMPRSS2-ERG fusion positive (TMPRSS2-ERG+) samples, like that shown relative to PDE4D7 score [Fig. 1C].

## Discussion & Conclusion

Our data shows that PDE4D is the only protein coding gene exhibiting an identical expression pattern to PDE4D7. Böttcher *et al*. (2016) noted that long isoforms of the PDE4D family are significantly downregulated during disease progression. However, as PDE4D5 and PDE4D9 are downregulated in both AS PCa and CRPC tissue whereas PDE4D7 is only diminished in CRPC, it represents a more specific measurement of disease progression (7).

Other RNAs mapped to the PDE4D coding region also exhibiting PDE4D7-like downregulation are pseudogenes (SETP21, RNU6-806P), nucleotide sequences (AC034234.1 and AC109486.1) and long non-coding (lnc) RNAs [prostrate androgen-regulated transcript-1 (PART1)]. Aside from PART1, no function has been ascribed to these genes, however PART1 has been extensively studied in cancer progression. PART1 is predominantly expressed in the prostate, relies on androgens for transcriptional regulation in PCa and is upregulated in PCa tissue (9,10). Interestingly, whilst PART1 is regulated via androgen signalling, PDE4D7 transcription is not, despite the PDE4D7 5’UTR coding region overlapping with antisense PART1 (4). Henderson *et al*. (2014) reported that PART1 and PDE4D7 exhibit positive correlation in mRNA expression within PCa cell lines and xenografts, which our results further confirm. However, we are the first to report the similarity in expression amongst various genes located within the PDE4D7 coding region of Chr5q12.

miR-582, which has also been implicated in PCa metastasis, showed correlation with PDE4D7 expression however the association was less robust than that of PART1. Similarly to PDE4D7, downregulated expression of miR-582-3p and miR-582-5p is associated with an advanced PCa phenotype with diminished expression observed in bone metastatic PCa tissues resulting in reduced bone metastasis-free survival (11). It is notable that although miR-582 expression correlated with PDE4D7 in PCa progression, there is high variability among patient samples, with many exhibiting no expression at all.

Correlation between PDE4D7 level and TMPRSS2-ERG status has previously been identified, with upregulated PDE4D7 in TMPRSS2-ERG+ tumours and low-grade PCa (8). Given that the PDE4D7 gene contains an ERG binding site, it was speculated that PDE4D7 may be regulated via ERG transcription. Our data further support this, with TMPRSS2-ERG-samples indicating downregulation of the analysed genes akin to those identified upon stratification by PDE4D7 score.

In summation, our data identifies the downregulation of an entire chromosomal region of Chr5q12 mapping to the PDE4D7 domain throughout PCa progression. This pinpoints not only driver genes but also passenger genes which may be a feature of carcinogenesis. PDE4D7 and PART1 have already been proposed as separate biomarkers for PCa (1,10), however in combination or alongside those identified here with correlative expression, could further enhance the specificity of prognostic approaches, improving diagnostics and therapeutics.

## Ethics

The local Institutional Review Boards approved the collection of patient tissue for clinical research.

